# Distribution maps of twenty-four Mediterranean and European ecologically and economically important forest tree species compiled from historical data collections

**DOI:** 10.1101/464834

**Authors:** Nadine Wazen, Valentina Garavaglia, Nicolas Picard, Christophe Besacier, Bruno Fady

**Affiliations:** Institut National de la Recherche Agronomique (INRA), Unité de recherches “Ecologie des Forêts Méditerranéennes, UR629 (URFM), Avignon, France.; Food and Agriculture Organization of the United Nations, Rome, Italy

**Author notes:** **Key message** Species distribution maps are often lacking for scientific investigation and strategic management planning at international level. Here, we present the range-wide, natural distribution maps of twenty-four Mediterranean and European forest-tree species of key ecological and economic importance in the Mediterranean. Dataset access: https://doi.org/10.5281/zenodo.1308577. Associated metadata access: http://www.fao.org/geonetwork/srv/en/metadata.show?id=56996.

## Background

Information on species geographic distribution is a strategic scientific resource for many research, innovation and development purposes, such as: biodiversity assessment, habitat and species management, restoration and conservation as well as for predicting the effects of global environmental change on ecosystems, species and populations and their genetic resources (Fady et al. 2016; Franklin 2009; Noce et al. 2016; Sinclair et al. 2010).

Maps are one of the ways information on geographic distribution can best be summarized and used (Pedrotti 2013). However, species distribution maps are often lacking or not made readily available for scientific investigation and strategic management planning at international level. In Europe, EUFORGEN, the program for genetic resource conservation (http://www.euforgen.org/), as well as the Joint Research Centre (JRC) of the European Union (San-Miguel-Ayanz et al. 2016) have made distribution maps available for many European species. Unfortunately, they address too rarely species of importance for Mediterranean countries outside of Europe. With climate change recently added to the long list of human impacts on Mediterranean forests, threats on their exceptionally rich biodiversity and on the livelihood of local communities are likely to increase (Médail and Quézel 1999; FAO 2014, 2015). Identifying valuable genetic resources and habitats to preserve is of the utmost importance in this context and the use of information on geographic distribution is a key step in this process.

Here, we present range wide, natural distribution maps of twenty-four Mediterranean and European forest tree species of key ecological and economic importance for countries of the Mediterranean Basin.

## Methods

### Sources of data

The 24 forest tree species (Table 1) were selected for their high economic and ecological importance by a panel of forestry experts from Algeria, Lebanon, Morocco, Tunisia and Turkey (see Acknowledgments section).

**Table 1.** List of the 24 forest tree species of high ecological and economic importance in the Mediterranean which were mapped.

|  |  |
| --- | --- |
| *Acer hyrcanum* subsp. *tauricum* (Boiss. & Balansa) Yalt. | *Arbutus unedo* L. |
| *Cedrus atlantica* (Endl.) Manetti ex Carriere | *Cedrus libani* A. Rich. |
| *Chamaerops humilis* L. | *Ilex aquifolium* L. |
| *Juniperus drupacea* Labill. | *Juniperus excelsa* M.-Bieb. |
| *Juniperus oxycedrus* L. | *Juniperus phoenicea* L. |
| *Laurus nobilis* L. | *Pinus brutia* Ten. |
| *Pinus halepensis* Mill. | *Pinus nigra* J.F. Arnold |
| *Pinus pinea* L. | *Pistacia lentiscus* L. |
| *Platanus orientalis* L. | *Quercus canariensis* Willd. |
| *Quercus cerris* L. | *Quercus coccifera* L. |
| *Quercus ilex* L. | *Quercus suber* L. |
| *Taxus baccata* L. | *Tetraclinis articulata* (Vahl) Mast. |

Data on the geographic distribution of the 24 species were compiled from the European Forest Genetic Resources Programme database (EUFORGEN, http://www.euforgen.org/), from published floras, from scientific publications containing syntheses of compiled data, and from the database of the Centre for Applied Research in Agroforestry Development (IDAF, Spain). The EUFORGEN data consisted of shapefiles defining distribution areas and the IDAF data consisted of geographical points of occurrence. Floras and other publications provided most of the distribution maps we used, in various image formats (Aytar et al. 2011; Bohbot et al. 2005; de Bolòs & Vigo 1984-2001; Boulos 1999; Browičz & Zielinski 1982; Committee for Mapping the Flora of Europe 1972-2013; Davis 1965-1988; Emberger 1939; FAO 2012; Fennane 1987, 1999; Gounot & Schoenenberger 1966, 1967; Lebanese Ministry of Agriculture 1965; Médail 2012; Quézel & Médail 2003; Quézel & Santa 1962-1963; Turkish Ministry of Forests and Water Affairs 2013; Yaltırık 1984).

Due to the difficulty of gaining access rights to country-level institutional databases, raw data from national forest inventories were not used, with the exception of the data from the Algerian national forest inventory that were used to locally refine the geographical distribution of some species. Although scientific publications in the field of ecology and forestry may report on the occurrence of the targeted tree species, no systematic review of these publications was made due to the high number of references (e.g. a search in the Web of Science using *Pinus halepensis* as keyword yielded over 1500 journal references) and to their redundancy with existing synthesis.

Information on the countries where the species are considered as native was collected from four databases: the Catalogue of Life (http://www.catalogueoflife.org/), the EURO+MED Plantbase (http://ww2.bgbm.org/EuroPlusMed/query.asp), the Kew World Checklist (http://apps.kew.org/wcsp/home.do), and the Med-Checklist (http://ww2.bgbm.org/mcl/query.asp). Country definition in these databases does not necessary match the administrative boundaries of countries but can correspond to biogeographically important regions within countries or to groups of countries. In the Med-Checklist for example, Italy as a country was split into Sicily, Sardinia and continental Italy while Lebanon and Syria, on the contrary, were grouped together to report on native species.

### Building the distribution maps

Maps were produced using existing digital distribution maps and digitizing by eye from published maps, hand-drawn maps on paper and other compilations. Georeferencing procedures were done using well-identified geographical reference points, and maps were then digitized into shapefiles. All operations were performed using QGIS 2.0.1, in particular QGIS Georeferencer plugin for georeferencing. In total, more than 100 maps were digitized requiring more than 18,000 entries (points or polygons) to be created. The countries of native distribution for each species were mapped using the FAO Global Administrative Unit Layers 2012-2103 shapefiles (http://www.fao.org/geonetwork/srv/en/main.home) to generate country boundaries.

Information on countries of native distribution was cross-checked with areas of distribution and points of occurrence to detect inconsistencies. A difficulty regarding points of occurrence is that the status of the presence of the species as a native species or as resulting from an introduction (botanic garden, plantation, etc.) was often not documented. This is one of the reasons we did not use the occurrence data of the Global Biodiversity Information Facility database (GBIF, http://www.gbif.org). When inconsistencies were found, countries of native distribution were checked with botanist experts and corrections incorporated. When uncertainties remained on how to solve these inconsistencies, no correction was made.

Points of occurrence were turned into areas of distribution using alpha-shapes. Alpha-shapes extend the concept of convex hull to recover the shape of a point cloud allowing this shape to be non-convex, multi-part, or with holes inside (Pateiro-López & Rodríguez-Casal 2010, Capinha & Pateiro-López 2014). Alpha-shapes depend on a parameter α that defines how wide or narrow the hull is. Specifically, two points will be allocated to the alpha-shape if there exists a circle of radius α with both points on its boundary, and which contains no other points. Individual points may remain outside of the alpha-shape as isolated points if they are too far away from other points. Polygons of occurrence of the species were computed in the Universal Transverse Mercator (UTM) coordinate system for geographical coordinates and using α = 50 km.

## Access to data and metadata description

For each species, known distribution localities were compiled into a vector shapefile (ESRI format) with point geometry. All species except *Cedrus atlantica* and *Chamaerops humilis* have such a shapefile with point locations. Countries (or regions within countries) or native distribution areas were compiled into a vector shapefile with polygon geometry. All species have a shapefile of country distribution. The presumed areas of native distribution were compiled into a vector shapefile with polygon geometry. All species except *Acer hyrcanum* subsp. *tauricolum* and *Juniperus drupacea* have such a shapefile of distribution area. We produced 68 shapefiles totalling 31 249 geometric objects (points or polygons). Each shapefile has a table of attributes that provides details on: i) scientific name, ii) common name, iii) country where the species have been reported, iv) data source, v) additional comments (Table 2). All maps are available in electronic format as shapefiles at: https://doi.org/10.5281/zenodo.1308577 (Wazen et al. 2018). Associated metadata are available at: http://www.fao.org/geonetwork/srv/en/metadata.show?id=56996. Maps (pdf format) showing the three geometries (localities, countries, area) of the species distribution are available as figures in this article and can be downloaded at http://www.fao.org/forestry/89249/en/.

**Table 2.** Table of attributes of the shapefiles giving the distribution range of 24 key Mediterranean and European tree species.

| Attribute | Description | Comments |
| --- | --- | --- |
| Country | Country where polygon or point is located | Not all fields are filled for polygons since some cover more than one country |
| Sp1_Sci_Na | Scientific name of Species 1 |  |
| Sp1_Com_Na | Common name of Species 1 |  |
| Sp2_Sci_Na | Scientific name of Species 2 (when information is available) | Polygons only: In some references, the area of occurrence is described with multiple species. In some cases the species of concern are mentioned as the 2nd, 3rd or 4th species; other times 2 or more species of concern appear in the same polygon and therefore are mentioned as 1st and/or 2nd and/or 3rd and/or 4th species |
| Sp3_Sci_Na | Scientific name of Species 3 (when information is available) |  |
| Sp4_Sci_Na | Scientific name of Species 4 (when information is available) |  |
| Source_Dat | The original map or reference used to get the data | For published data, the attribute consists of the author, date and title as listed in the References |
| Comments | Any relevant comment regarding the data |  |
| code_map | the code on the original map that refers to the specific species (when applicable) |  |

## Technical validation

Polygon geometries gave information on the presence (inside the polygon) and absence (outside) of the species. Depending on the data source, they were provided with different levels of spatial detail (Figure 1). Point geometries either informed on the punctual presence of the species (but not its absence elsewhere), or on its presence in cells of a grid system when points were arranged on a grid (Figure 1).

**Figure 1.**
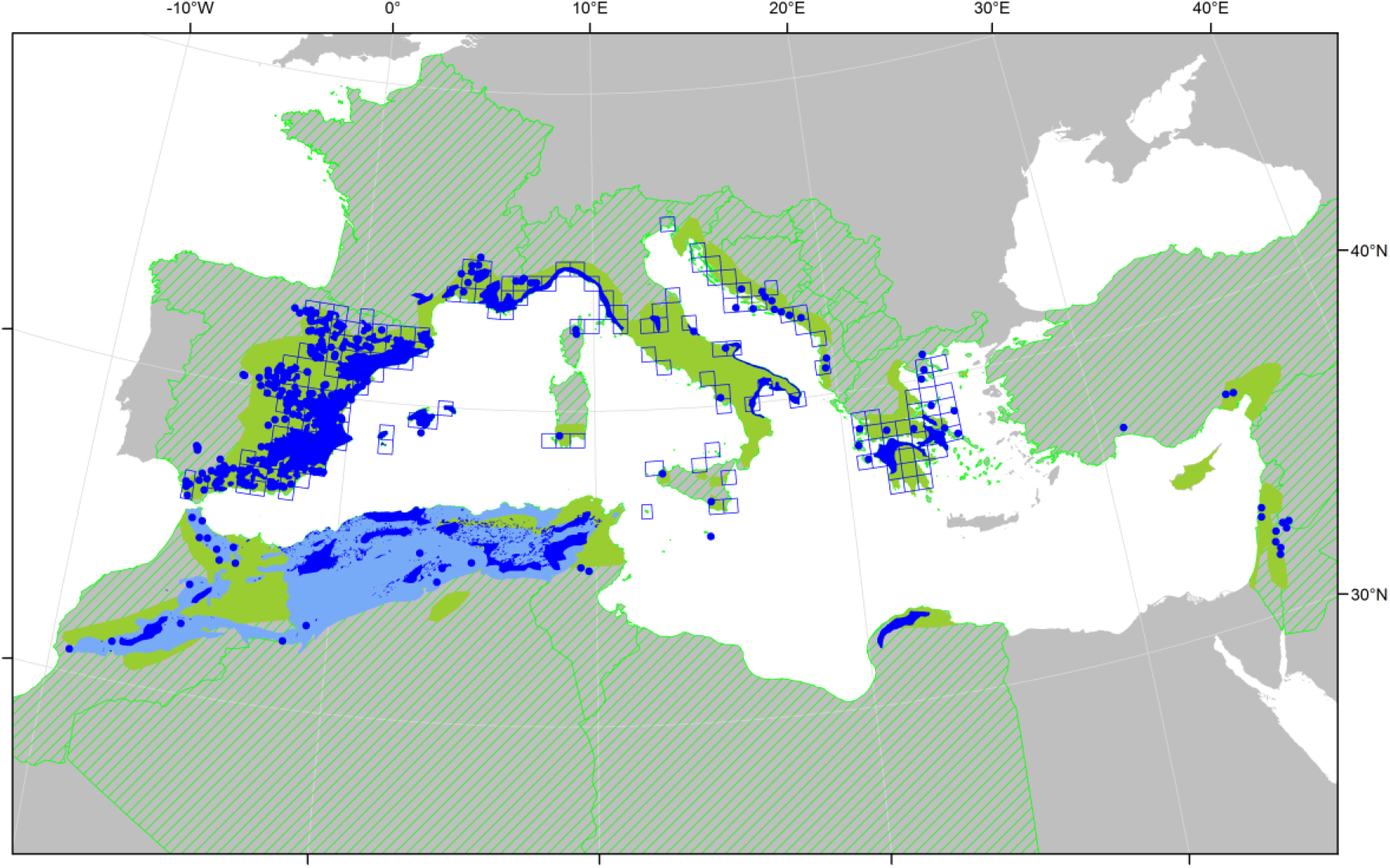
Distribution map of *Pinus halepensis* showing the different types of data collected: countries of native distribution (hatched green polygons); areas of native distribution with high (blue filled polygons), medium (light blue polygons), or low (green polygons) level of spatial details; points of occurrence represented either as points (blue points) or as the cells of the Common European Chorological Grid Reference System where the points were found (blue unfilled polygons).

These heterogeneous data sources were combined for each species to create synthetic species distribution maps, with the different levels of details from the different data sources possibly resulting in polygon overlaps and finer details being masked. When aggregating neighbouring points into polygons, we checked that the projection system had little influence on the outcome. Shorter values than 50 km for α were not appropriate because points of occurrence were sometimes distributed along a grid and α < 50 km failed to connect the points across this grid. Longer values than 50 km for α tended to create unrealistically wide distribution areas. As an example, Figure 1 shows the distribution area computed as the alpha-shape of points of occurrence of *Acer hyrcanum* subsp. *tauricolum* using α values ranging from 30 to 100 km. All computations were made using the R statistical environment (www.r-project.org) and the alphahull package to compute alpha-shapes (Pateiro-López & Rodríguez-Casal 2010). The maps where the point geometry of localities were converted into polygons using the alpha-shapes and merged with areas, are shown in Figure 2.

**Figure 2.**
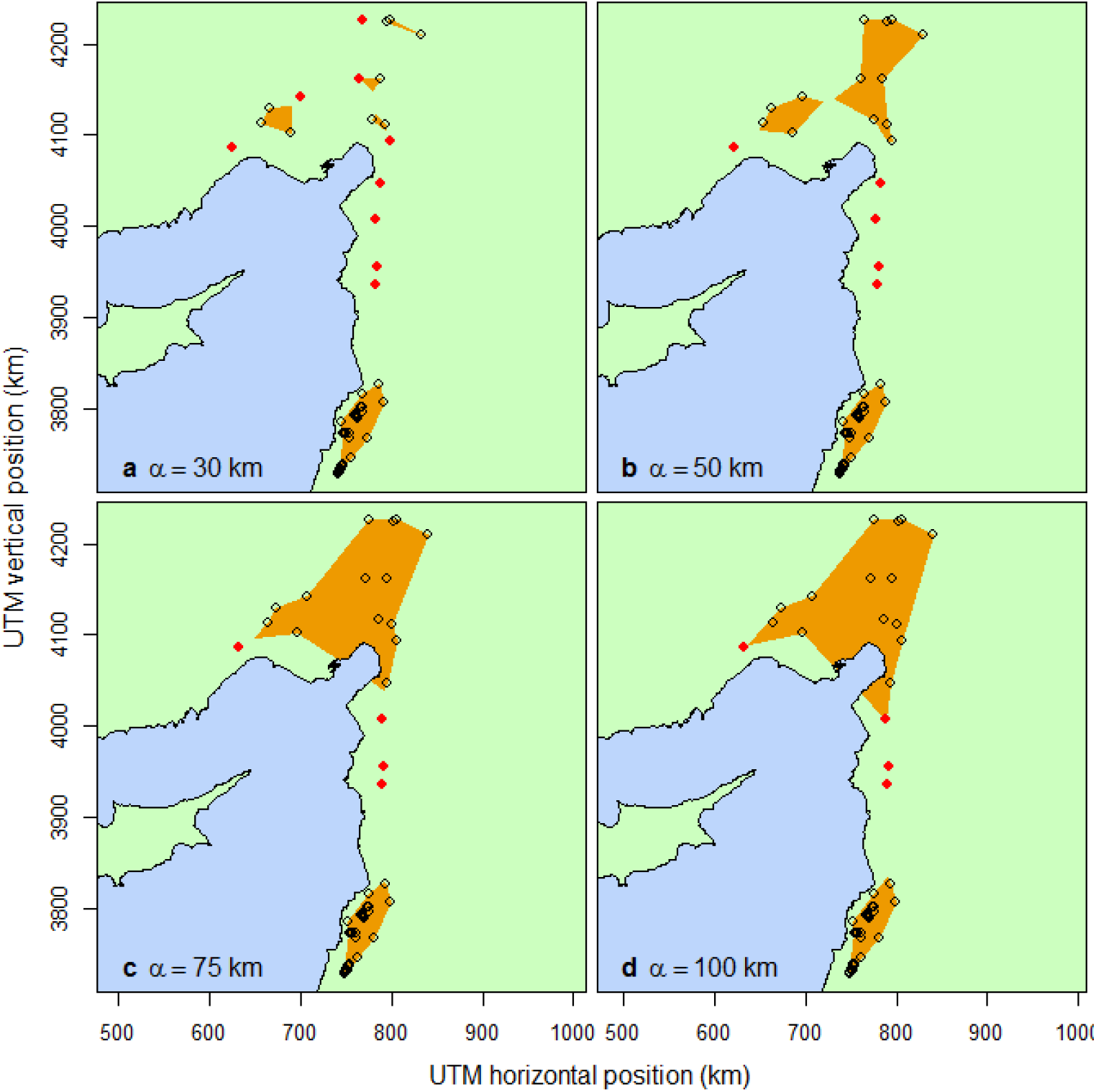
Alpha-shape (orange polygon) of points of occurrence of *Acer hyrcanum subsp*. tauricolum using α values ranging from 30 to 100 km. Points of occurrence included in the alpha-shape are shown as black dots while points of occurrence that remain isolated points (outside the alpha-shape) are shown as red dots. The UTM zone is 36S.

## Reuse potential and limits

We generated a set of distribution maps for 24 forest tree species (Figure 3), key for Mediterranean forestry, which we consider as a strategic resource for both science and management. These maps are general range maps, created from the compilation of multiple sources of information (mostly published floras and chorological maps) with different levels of accuracy, where in some cases the most recent available data was decades old. The alpha-shape procedure used to turn occurrence data points into polygons purposefully degraded precise information, with the possibility that polygons may overlap and further blur local details. However, the maps are meant to be accurate only at the level of the entire distribution range of the species or at country level and not at finer spatial scales.

**Figure 3.**
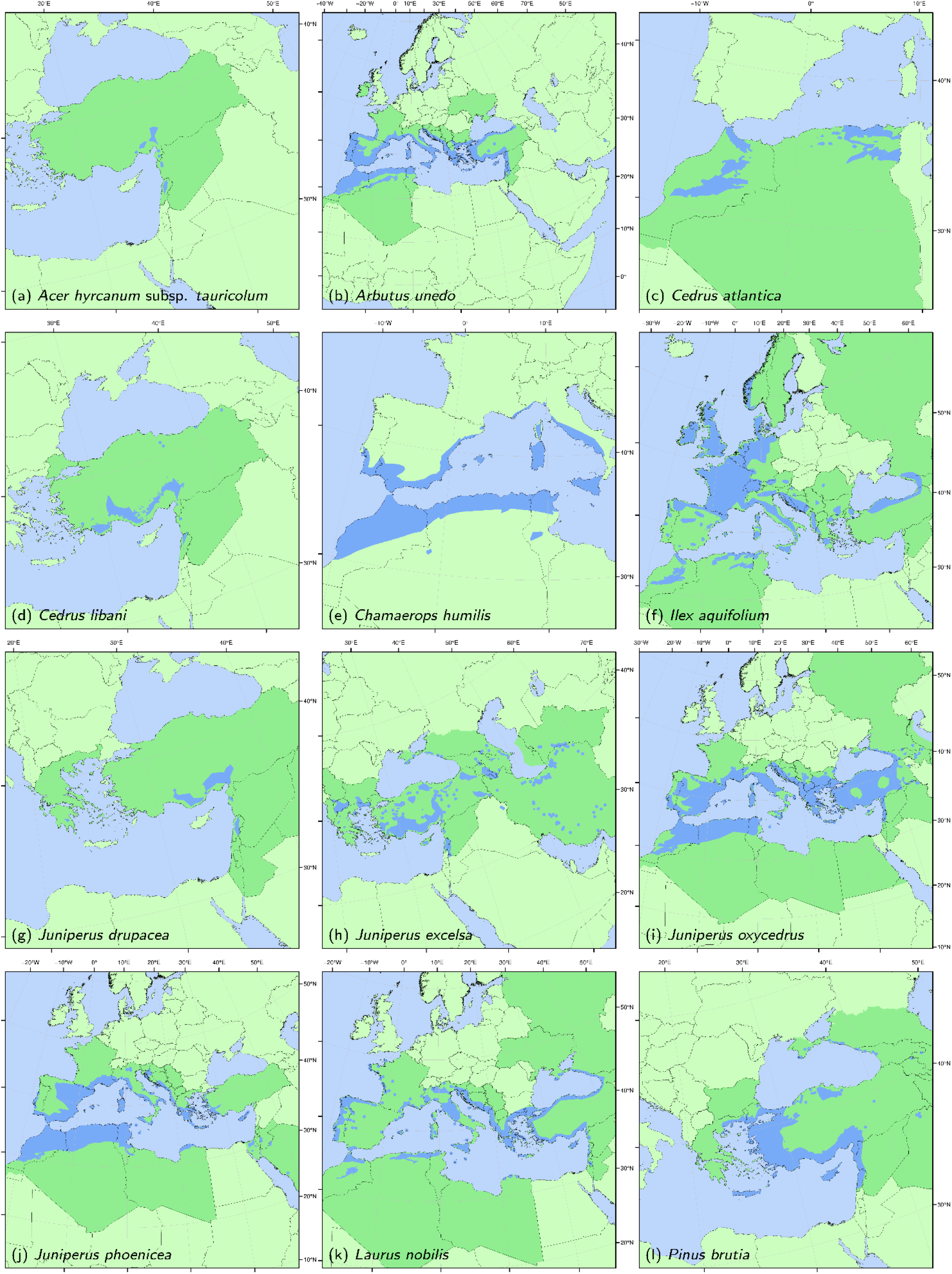
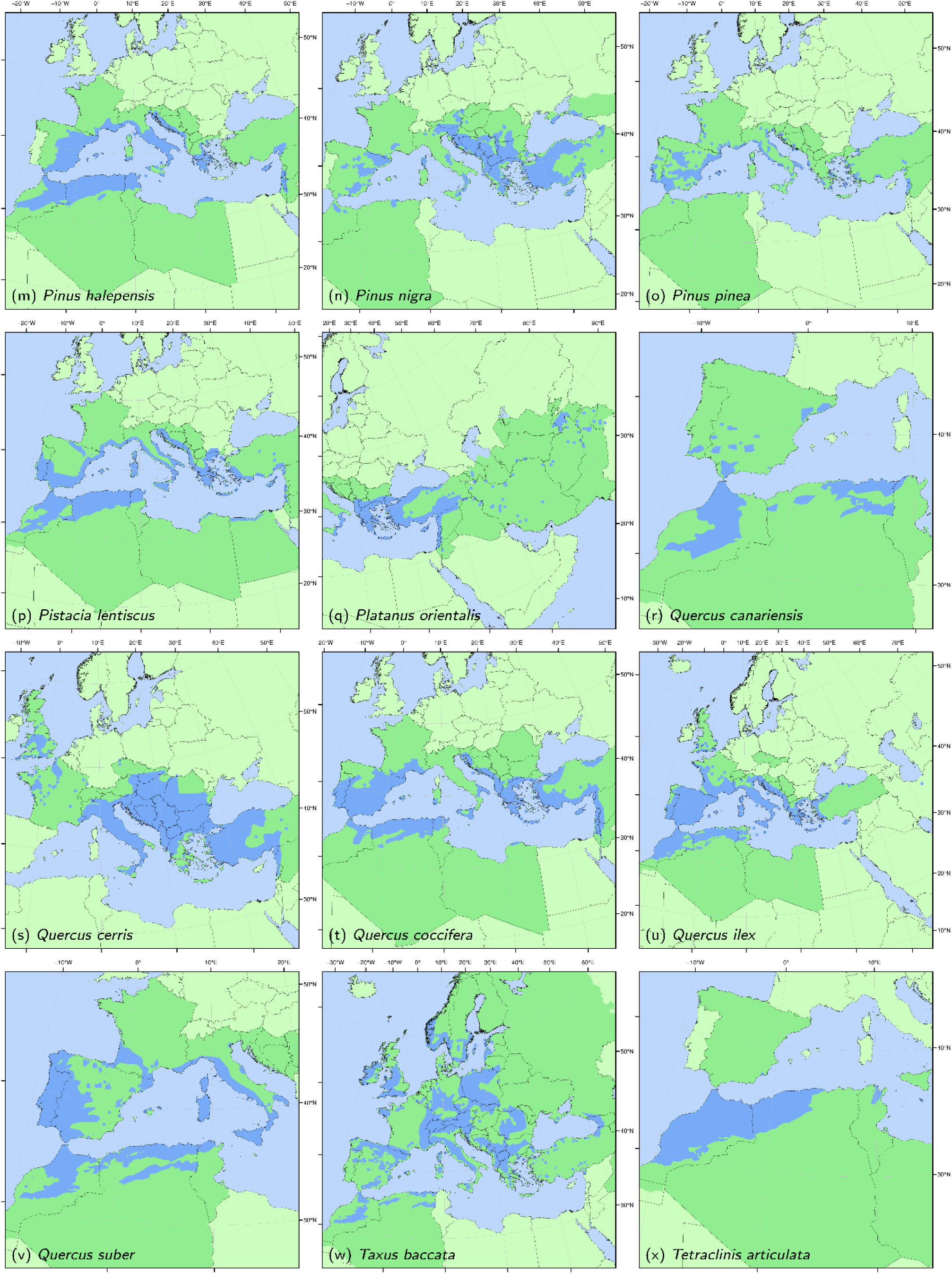
Distribution maps of 24 key Mediterranean and European forest tree species based on the compilation of published data. Green shows the countries (or regions within countries) of native distribution of the species. Blue shows the presumed area of native distribution, where localities of known distribution have been merged into an area using alpha-shapes with α = 50 km.

We did not digitize all available published chorological maps and the works of Meusel & Jäger (1965-78-92), of Hultén & Fries (1986) or of Critchfield & Little (1966) for example, could be added to our dataset and might refine some distribution limits. The diversity of data resolutions used for drawing the maps, however, does not make it possible to downscale the information as is possible with resources made from occurrence data from inventories such as the European atlas of forest tree species (San-Miguel-Ayanz et al. 2016).

Our maps are not maps of exact occurrence at all spatial scales either. Geographic data compiled from published sources are provided both as a basis of knowledge and as a basis for further discussion and questioning on the distribution of the species. During our quality check procedure (see Assante et al. 2016 for a discussion on data quality), which included feedback from a panel of experts (see acknowledgements), we detected biases that we corrected while we decided to keep others. To give a few examples of such questionings raised by the maps, the presence of *Quercus ilex* in England may or may not be of native origin; the isolated localities of presence of *Arbutus unedo* in Iran may be questioned; or the occurrences of *Pinus halepensis* in Cyprus may not be real. These maps may also provide guidance on where further inventories should be conducted to clarify the distribution of the species. The eastern limit of the distribution map of *Cedrus atlantica* that coincides with the border between Algeria and Tunisia, for instance, calls for further investigation of this species in eastern Algeria.

Therefore, as more and more digital resources are made available and academic and citizen-science knowledge accumulates, we recommend that experts in forest tree species distribution indicate how ranges should be refined to adjust zones where the species are wrongly indicated as naturally occurring, by either manipulating and reposting the shapefiles or contacting the authors. Resources such as those of GBIF, properly documented, could be used for such a purpose.

## Contribution of the co-authors

Nadine Wazen: compilation of the data, digitization, data quality control and analysis, writing the paper.

Valentina Garavaglia: digitization, data quality control and analysis, writing the paper Nicolas Picard: digitization, data quality control and analysis, data analysis, writing the paper Christophe Besacier: coordination of the research project

Bruno Fady: coordination of the research project, supervising the work, writing the paper.

## Acknowledgements

Partners of the FFEM project “Maximize the production of goods and services of Mediterranean forest ecosystems in the context of global changes” (project 2011/CZZ1695) selected the 24 species of interest analyzed here. A first version of the maps was presented at the XIV World Forestry Congress in Durban (South Africa) in 2015 and can be viewed at: http://foris.fao.org/wfc2015/api/file/55312a832e3571f323904b91/contents/297183f1-2b6c-4b9f-9e29-e67d92542323.pdf. The authors thank the project partners who provided information and data on the geographical distributions of the 24 tree species, in particular the General Directorate of Forests of Algeria, the Ministry of Agriculture of Lebanon, and the Centre for Applied Research in Agroforestry Development (IDAF, Spain). We also wish to thank: Prof. Frédéric Médail (Aix-Marseille University, France), Mr. Daniel Pavon (Aix-Marseille University), and Dr. Bouchra Douaihy (Lebanese University, Lebanon) for providing expert knowledge on species distribution and autochthony and pointing out discrepancies in some resources; Mr. Didier Betored (URFM) and Ms. Marianne Corréard (UEFM) at INRA Avignon (France), for providing help and support on geographic information system use and data management; Mr. Michele Bozzano (Bioversity International, Rome) for granting access to digitized EUFORGEN map data and providing support on methodology development.

